# Deep-learning based bioactive peptides generation and screening against Xanthine oxidase

**DOI:** 10.1101/2023.01.11.523536

**Authors:** Haiping Zhang, Konda Mani Saravanan, John Z.H. Zhang, Xuli Wu

**Author notes:** Corresponding to: Haiping Zhang and Xuli Wu.

## Abstract

In our previous work, we have developed LSTM_Pep to generate *de novo* potential active peptides by finetuning with known active peptides and developed DeepPep to effectively identify protein-peptide interaction. Here, we have combined LSTM_Pep and DeepPep to successfully obtained an active *de novo* peptide (ARG-ALA-PRO-GLU) of Xanthine oxidase (XOD) with IC50 value of 3.76mg/mL, and XOD inhibitory activity of 64.32%. Consistent with the experiment result, the peptide ARG-ALA-PRO-GLU has the highest DeepPep score, this strongly supports that we can generate *de novo* potential active peptides by finetune training LSTM_Pep over some known active peptides and identify those active peptides by DeepPep effectively. Our work sheds light on the development of deep learning-based methods and pipelines to effectively generate and obtain bioactive peptides with a specific therapeutic effect and showcases how artificial intelligence can help discover *de novo* bioactive peptides that can bind to a particular target.

## Introduction

Hyperuricemia is increasing prevalence in the globe, especially in Asian countries^1^. Xanthine oxidase (XOD) is a therapeutic target of hyperuricemia^2^. This enzyme was required to produce uric acid through the breakdown of purine nucleotides. Its inhibitors are used in the clinical treatment of hyperuricemia^3^. Xanthine oxidase inhibitors include purine analogues and others. Many natural products have been found to inhibit xanthine oxidase in vitro or in model animals, for instance, kaempferol, myricetin, and quercetin^4^.

Many peptides also can inhibit Xanthine oxidase and many of them are derived from protein hydrolysates of food ^5,6^. For instance, FPSV and FPFP from round scad (Decapterus maruadsi) protein hydrolysates exhibited XOD inhibitory activity^7^. A novel peptide named RDP1 (AAAAGAKAR) from the extract of shelled fruits of Oryza sativa could significantly reduce the serum uric acid and alleviate hyperuricemic in rats by inhibiting XOD, and RDP1 showed no toxicity in rats^8^.

Deep learning techniques have been widely applied in biological related fields. Recently, we have developed LSTM_pep for *de novo* potential bioactive peptide generation through finetune training over known active peptides, and DeepPep for prediction of whether given protein-peptide are binding^9^. The combination of LSTM_Pep and DeepPep can provide an effective way to generate and screen potential active peptides for given protein targets.

In this work, we successfully identified a *de novo* active peptide of OXD by using LSTM_pep and DeepPep together, and this active peptide (ARG-ALA-PRO-GLU) turn out to have the top DeepPep score.

## Method

### Screening *de novo* peptides for Xanthine oxidase (XOD) by LSTM_pep and DeepPep

To further demonstrate the strong power of the combination LSTM_Pep and DeepPep in identifying *de novo* peptides, we efficiently finetune training, generating, and screening to obtain potential *de novo* peptide inhibitors of XOD by LSTM_pep and DeepPep. We obtain 19 known active peptides of XOD from literature reports, shown in Table 1. We finetune training LSTM_Pep over these peptides and generate 5000 *de novo* peptides. After removing redundancy and peptides same as the training set, we finally obtained 821 peptides sequence. Then, we built 3D structure of the 821 peptides by TrRosetta software^10^.

**Table 1.**
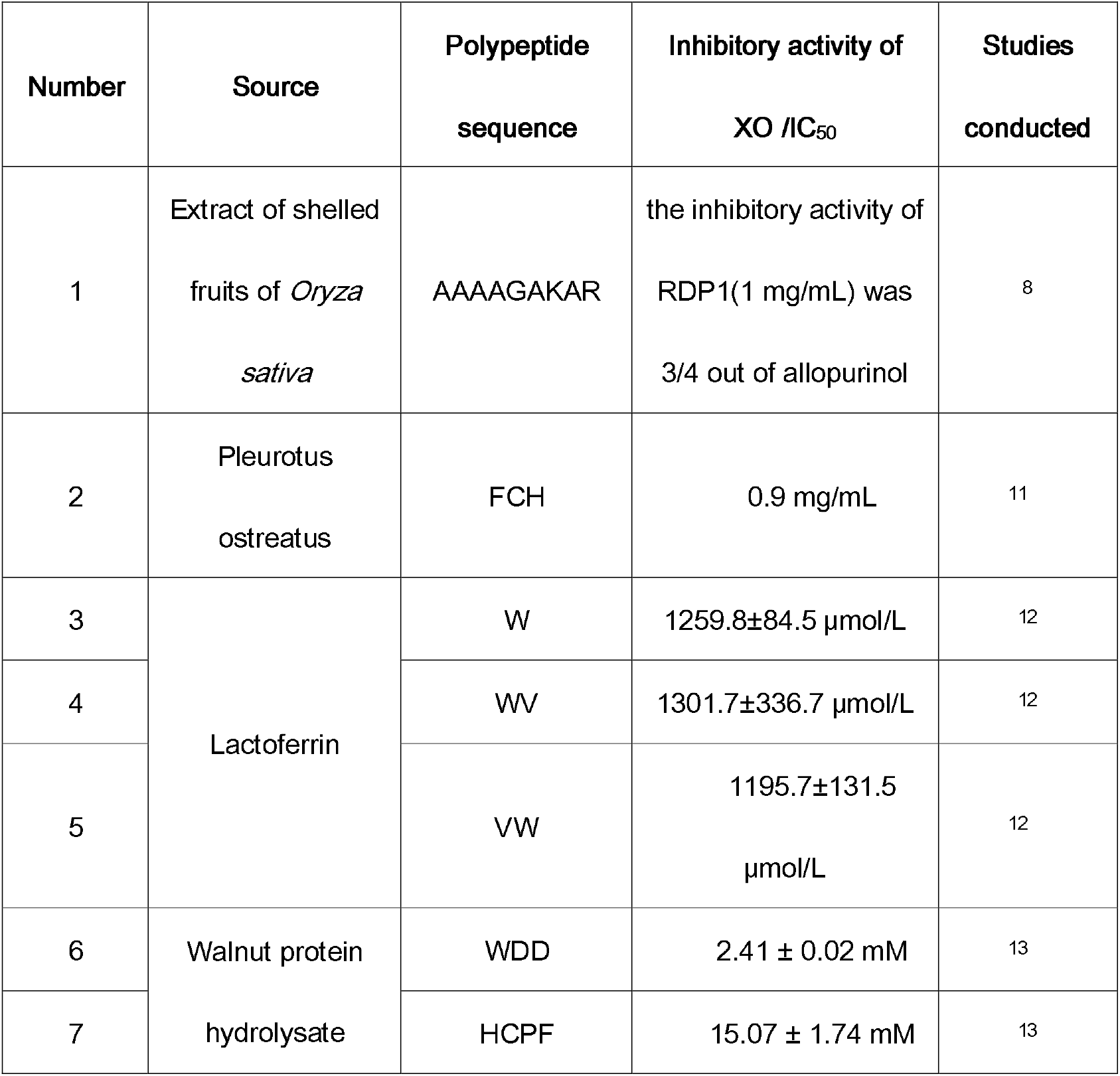
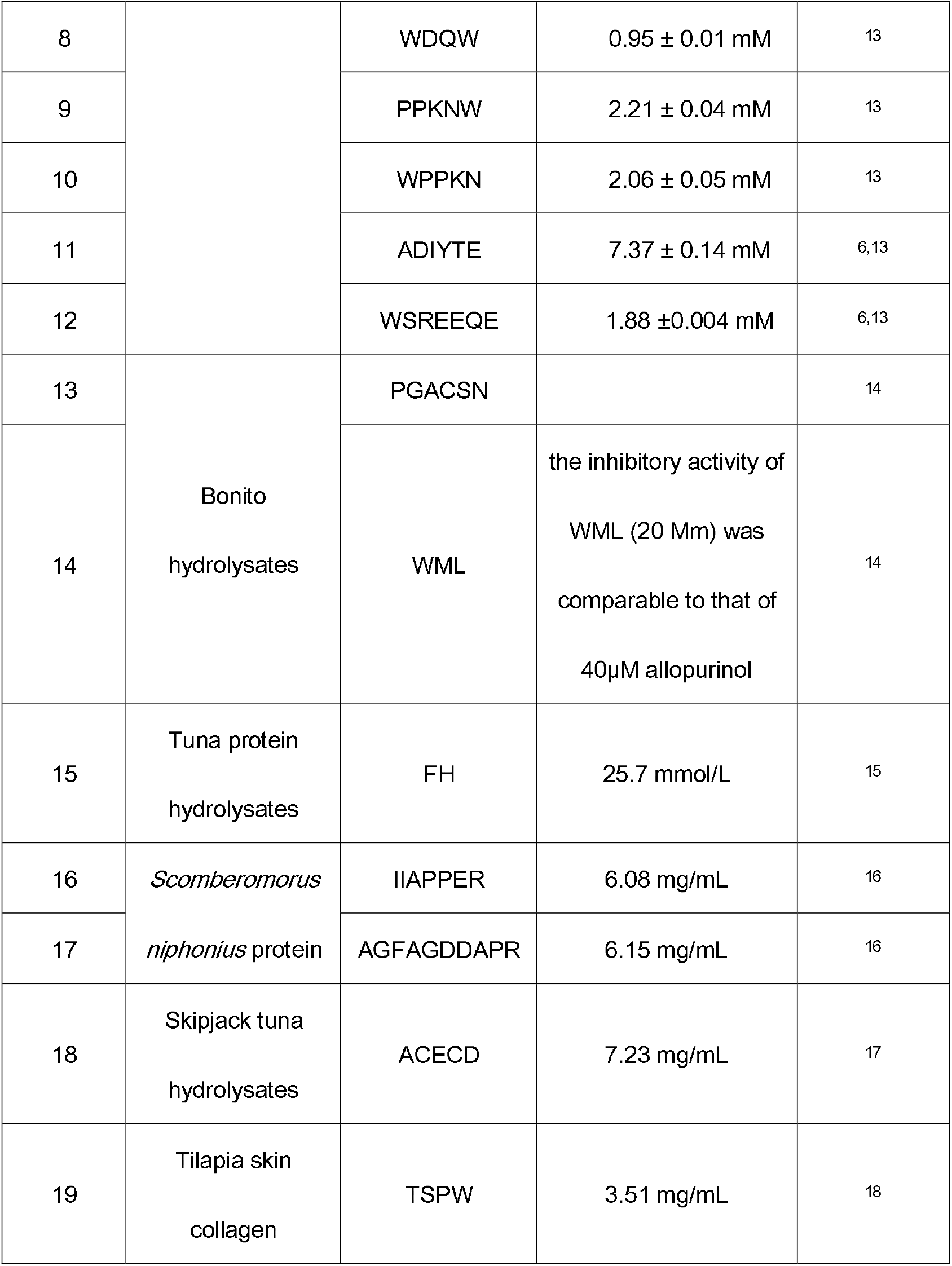
The information of 19 known peptide inhibitors of XOD. Their source, sequence, inhibitor activity/IC50, and reference were listed.

We obtain XOD’s ligand binding domain from the PDB database (PDBid 3nvy^19^, C chain, with cofactor MTE, MOS). The 821 obtained peptides’ 3D structure were then docked into the pocket of XOD with ZDOCK, the docked result contains 2000 binding conformation with docking scores. The top 200 docked conformations (by ZDOCK score) for each peptide-XOD pair were transferred into the figure-like matrix as input of the DeepPep. Finally, DeepPep was used to predict the binding possibility of these 200 conformations, and select the top prediction as the peptide-XOD pair score, and the corresponding conformation as the predicted binding pose. In such a way, we have scored and ranked all the peptide-XOD pairs. Finally, we obtain a candidate list with all *de novo* peptides that have DeepPep score larger than a cutoff value (*e*.*g*. 0.9), and five with top scores are selected for further experimental validation.

### Measurement of XOD inhibitory activity

XOD activity assay was conducted according to the method^20^ with slight modifications. Briefly, XOD (30 μL, 0.1 U/mL) was incubated with various amounts of the samples in phosphate buffer (0.05 mol/L, pH 7.5) at 37 °C for 5 min before the reaction. Then, the enzymatic reaction was started by adding 60µL of substrate xanthine (150 μM) to the mixture. The final absorbance measurement was read at 295 nm for calculating the enzyme activity. The XOD inhibitory activity in the enzymatic reaction was estimated as

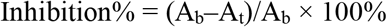

where A_b_ is the absorbance of the control and A_t_ is the absorbance of samples. The IC_50_ obtained from the inhibition curve represents the concentration of an inhibitor that inhibits 50% of XO activity.

### The predicted peptides for Xanthine oxidase (XOD) and experimental validation results

Hyperuricemia can lead to several serious diseases, including gout(a common form of inflammatory arthritis), cardiovascular disease, diabetes, and kidney disease^21^. In order to obtain *de novo* peptide inhibitors of XOD to combat hyperuricemia, we combine LSTM_Pep, DeepPep, and the experimental method together to generate and screen *de novo* XOD peptides efficiently in a step-by-step pipeline, shown in Figure 1.

**Figure 1.**
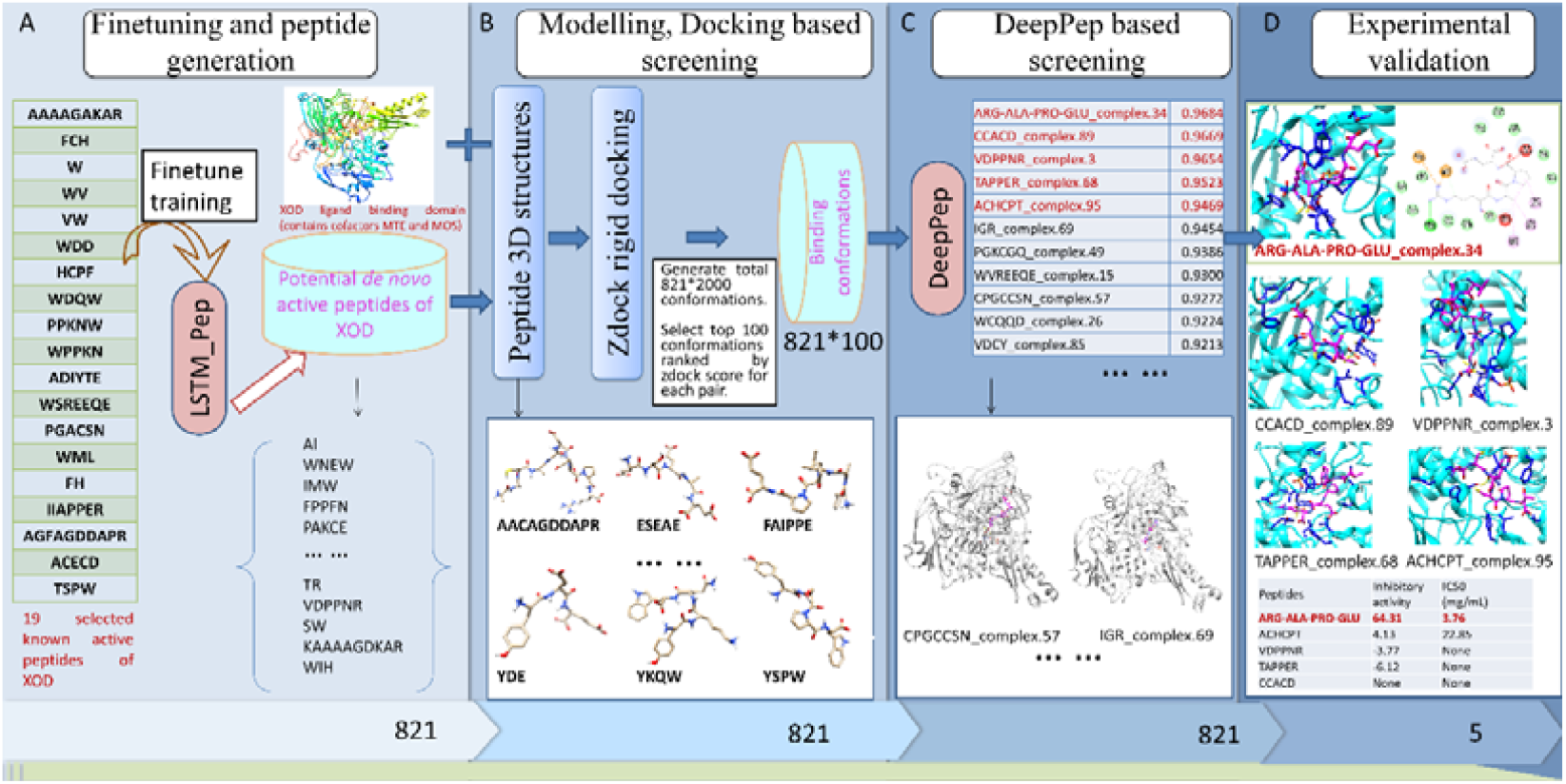
The workflow of combination LSTM_Pep and DeepPep in identifying *de novo* active peptides of XOD. Panel A, finetune training the LSTM_Pep model with 19 known active peptides of XOD, and generated 821 *de novo* peptides by finetune trained model; Panel B, building 3D structure for the generated *de novo* peptides by TrRosetta, then Zdock was used to dock the peptides into the pocket region of XOD (with cofactor MTE, MOS), for each docking, we selected top 100 conformations from total 2000 conformation; Panel C, DeepPep was used to predict the binding possibility of selected conformations, and score each XOD-peptide binding pair; Panel D, the top five predictions were selected for experimental validation.

we finetune training our LSTM_pep over 19 known active peptides of XOD, and using the obtained LSTM_Pep to generate 821 unique new potential Peptides, as shown in Figure 7A. The 821 unique new peptides were modeled into the 3D structures by TrRosseta software. Then, we docked those 3D peptide structure into XOD pocket, shown in Figure 7B. For each docking result, we obtain the top 100 zdock conformations for further DeepPep prediction. In other words, the DeepPep has carried a total of 82100 times predictions. Finally, we obtained a candidate list with DeepPep prediction score, among them 11 peptides have score values reached above 0.92, shown in Figure 3C. Finally, we select the top 5 candidates for experimental validation. The experimental results show that ARG-ALA-PRO-GLU has an inhibitory activity value of 64.31 and IC50 of 3.76 mg/mL, which is a very encouraging result, indicating ARG-ALA-PRO-GLU could be a good start for future modification to achieve even higher affinity.

## Conclusion

We find combined LSTM_Pep and DeepPep can generate and screen *de novo* active peptide very efficiently, and applying these two methods in target Xanthine oxidase led to the successfully discover an active *de novo* peptide (ARG-ALA-PRO-GLU). This work supports the practical usage of LSTM_Pep and DeepPep in accelerating the finding of active *de novo* peptides for a given therapeutic protein targets.

## Author contributions

H.Z. designed the study. H.Z., K.M.S, and X.W. performed computations and data analyses. All authors contributed to writing the manuscript. H.Z. and X.W. supervised the study. All authors read and approved the final manuscript.

